# Versatile and Cross-Species Alveolar Type II Cell Isolation Using Humanized Monoclonal Antibody Upifitamab

**DOI:** 10.64898/2026.11.10.765643

**Authors:** Michael T. Forrester, Shuyu Liu, Kyoshiro Tsuge, Rhianna Lee-Ferris, Krsna Rangarajan, R. Ian Cumming, Khaliun Enkhbayar, Jeremy M. Morowitz, Thomas J. Whitlow, Aleksandra Tata, Bin Li, Christine E. Eyler, Robert M. Tighe, Purushothama Rao Tata

## Abstract

Alveolar type 2 (AT2) cells are primary adult alveolar stem cells that maintain healthy alveoli. AT2 cell dysfunction is broadly linked to many lung pathologies from emphysema to fibrosis. Isolation of AT2 cells with high purity is therefore central to diagnostic studies, proteomics, transcriptomics and ex vivo experiments that can lead to future AT2 cellular therapies. Current markers for AT2 isolation exhibit various limitations from species-restrictions (e.g., mouse vs human) to limited AT2 specificity, which creates challenges for flow cytometric gating/sorting and application to magnetic-activated cell isolation. While underrepresented in the literature, these bottlenecks have made AT2 cell isolation inherently difficult and slowed AT2 cell research. Here we leverage surface protein-restricted transcriptomics to identify the sodium phosphate transporter NaPi2b (SLC34A2) as a highly specific AT2 cell marker optimally detected with the humanized monoclonal antibody upifitamab. Upifitamab-based labeling of AT2 cells is compatible with flow cytometric or magnetic-activated AT2 isolation and applies to both mouse and human AT2 cells, thus streamlining protocols and circumventing species-specific nuances. AT2 cells obtained with upifitamab can be cultured as organoids, in monolayers or differentiated to AT1 cells. This simplified AT2 isolation approach should overcome several current barriers while facilitating *ex vivo* studies on alveolar biology and related therapeutic developments.

## INTRODUCTION

Alveolar type 2 (AT2) cells are cuboidal epithelial stem cells that perform a wide range of homeostatic functions including surfactant synthesis^1^, immunologic coordination^2,3^ and replenishment of gas-exchanging alveolar type 1 (AT1) cells during injury^4–6^. Dysfunction of AT2 cells is linked to a wide range of conditions from emphysema^7^ to interstitial lung diseases such as idiopathic pulmonary fibrosis (IPF)^8^. Considering these integral roles in physiology and pathophysiology, AT2 organoids have received significant attention over the past decade^9,10^ as a culture method that recapitulates critical elements of alveolar biology.

Current methods for AT2 isolation generally utilize fluorescence assisted cell sorting (FACS) coupled to immunostaining for epithelial cell adhesion molecule (EpCAM, CD326). However, EpCAM is broadly expressed across the epithelium and has been used to isolate human bronchial epithelial cells^11^, highlighting its insufficient specificity for AT2 cells. This shortcoming of EpCAM to differentiate epithelial subtypes has been partially addressed through the addition of IA/IE (MHCII) staining^12^ or pH-dependent fluorophore uptake of Lysotracker dyes^13^. A major limitation for MHCII is its prevalence in immune cells and lack of expression on human AT2 cells (i.e., mouse-restricted). Similarly, Lysotracker staining also requires CD45 negative gating (due to high immune cell uptake), has shown wide variability across human lung specimens^14^, is not amenable to magnetic-activated cell sorting (MACS) and does not survive cell fixation. Due to these technical limitations, transgenic mice expressing AT2-restricted fluorophores continue to be frequently employed for AT2 cell isolation by FACS. However, this approach necessitates dedicated lineage-labelled transgenic mice, cannot be applied to human tissues and is not compatible with MACS.

Another limitation of current methods is that minor amounts of contaminating airway cells can rapidly outgrow AT2 cells in culture^15^. An example are distal airway secretory cells, which can be distinguished from AT2 cells by their CD66 positivity^16,17^. However, CD66 is poorly conserved in mouse and gating out CD66+ cells in human lung has shown marginal benefit in terms of increasing AT2 cell purity^14^.

Isolation of human AT2 cell isolation was enhanced by generating hybridoma clones displaying AT2-specific immunoreactivity^18^. This so-called antibody clone (HTII-280) has allowed for MACS isolation of human AT2 cells using a single antibody enrichment step, facilitating many AT2-oriented studies. This antibody may recognize a glycosylated region of human MUC1^19^, which is itself broadly expressed on lung epithelium and not particularly specific to AT2 cells. Ultimately, HTII-280 has several important limitations including 1) an incompletely characterized epitope, 2) lack of applicability to mouse AT2 cells and 3) issues inherent to IgM antibodies such as aggregation, polyvalency, steric bulk and complement activation. To address these methodological shortcomings and nuances between human vs mouse AT2 cells, we sought to identify a surface epitope-antibody pair that could allow for simple cross-species AT2 cell isolation by multiple methods (e.g., MACS, FACS).

## METHODS

### Transcriptomic sub-analyses of surface associated proteins

Candidates were restricted to proteins with potential for cell-surface expression, defined by the SURFY *in silico* human surfaceome^20^. This list has 2,886 predicted surface proteins corresponding to 2,799 unique gene symbols. For mouse datasets, human genes were mapped to mouse orthologs with the Mouse Genome Informatics (MGI) human–mouse homology table (HOM_MouseHumanSequence.rpt)^21^ covering 2,554 genes. The remaining 245 genes were matched by gene symbol.

Four human and five mouse lung scRNA-seq datasets were analyzed. Human data were the Human Lung Cell Atlas (HLCA) core^22^, the lung portion of Tabula Sapiens^23^, the Krasnow Lab lung atlas^24^ and a bronchopulmonary dysplasia (BPD) dataset (CELLxGENE collection 3a5dbf8a-9b3e-4309-b4c5-d8a024f83734). Mouse datasets include the postnatal (P7, 2-week and adult) cells of the LungMAP developmental atlas^25^, a bleomycin injury time course with PBS controls (GSE141259)^26^, an aging lung atlas (GSE124872)^27^, and a neonatal normoxia/hyperoxia dataset (GSE151974)^28^. The HLCA, Tabula Sapiens, Krasnow Lab, BPD and LungMAP data were obtained from CZ CELLxGENE Census (release 2025-11-08)^29^.

Cell-type labels were used as provided by each source, with no re-clustering or re-annotation. AT2 cells were those labelled “pulmonary alveolar type 2 cell” (Cell Ontology; Census datasets), “AT2 cells” (GSE141259), “Type_2_pneumocytes” (GSE124872), or “AT2 1” and “AT2 2” combined (GSE151974). All other annotated cells formed the comparison group. Expression was library-size normalized by dividing each cell’s counts by total counts.

Three metrics were computed for each surface gene in each dataset. (i) AUROC: the area under the receiver operating characteristic curve, using the gene’s expression as the only classifier of AT2 versus all other cells. This measures how well the gene separates AT2 cells from everything else. 0.5 = no separation and 1 = perfect separation. (ii) Tau (τ): a tissue-specificity index^30^ computed over the mean expression of each annotated cell type as τ = Σ_i_(1 − *x*_i_/*x*max)/(*n* − 1). τ ranges from 0 (uniform expression) to 1 (expression in a single cell type). (iii) log2 fold change: log2 of the mean AT2 expression divided by the median of the other cell types. The log2 fold change was ranked only among genes detected in ≥20% of AT2 cells due to ratio instability for sparsely detected genes. Genes were ranked by AUROC among all detected surface genes and by log2 fold change among the ≥20%-detected subset. Results are shown as scatter plots of τ against AUROC, plotted on a log(1 − AUROC) scale to resolve values near 1. All analyses used Python 3.12 with cellxgene-census 1.18.0, anndata 0.12.18, NumPy, pandas, SciPy, scikit-learn 1.9.0 and Matplotlib.

### Cloning and lentiviral transduction of human Cyclophilin A (CyA) or mouse Slc34a2 (NaPi2b) in HEK293T cells

ORFs for human Cyclophilin A (gene *PPIA*; UniProt P62937) and mouse *Slc34a2* (UniProt Q9DBP0) were synthesized by Twist Biosciences, digested with XbaI and EcoRI then purified by gel extraction. For Slc34a2, a C-terminal HA tag was included in the construct design. Digested ORFs were ligated into pFW2-MCS, which is an in-house lentiviral vector utilizing a full EF1 promoter and T2A-Hyg for hygromycin resistance. All plasmids were confirmed by whole plasmid sequencing (Genewiz / Azenta).

Lentiviruses were produced in 10 cm dishes of HEK293T cells cultured in Dublecco’s Modified Eagle Medium (DMEM) with 10% FBS, 10 U/mL Penicillin and 10 µg/mL Streptomycin. At 80-90% density, cells were transfected with 14 µg pFW-CyA or pFW-Slc34a2-HA, 12 µg psPAX2 and 4 µg pMD2.G using a 4:1 ratio of 40 kD Linear PEI : DNA. Media was changed at 24h, then collected at 48, 72 and 96 h. The 30 mL of conditioned media was filtered through a 0.45 µm PES syringe filter followed by addition of 10 mL PBS containing 32% (w/v) PEG-6000 and 1.6 M NaCl. Concentration was performed at 4C for 24h then collected by centrifugation at 2000g x 1h. Pellets were resuspended in a 50^th^ volume (0.6 mL) and frozen at −80C (100 µL x6 aliquots). For transduction, 100 µL of concentrated virus was added to one well of a 12 well dish containing HEK293T cells at 40-50% confluence. On day 3, hygromycin was added at 250 µg/mL and selection was performed over 7-10 days. Cells were maintained in 200 µg/mL hygromycin during continued passages.

### Conjugation of isotype control IgGk and upifitamab with Alexa Fluor 647

All Zeba 7kDa column centrifugation steps performed at 1500g x 1 min in a fixed angle rotor with columns consistently oriented during each spin (by marking outer side of the column). Columns were equilibrated in 100 mM borate pH 8.5 x3 spins. To desalt and remove Tris, upifitamab or isotype control IgGk (100 µL, 2 mg/mL) was added to the top of each column. After 1 min, 20 µL of borate buffer was added as a stacker solution. Centrifuged. Flow through saved. Alexa Fluor 647-NHS (100 µg, Thermo) was resuspended in 80 µL of water final of 1 mM, then 18 µL immediately added directly to each antibody. Molar ratio calculations: Antibodies 1.66 mg/mL (9.6 µM), Alexa647-NHS 130 µM for molar excess of 13.6:1 dye:IgG. After 1h in the dark at room temp, 13 µL of 1 M Tris pH 8.0 was added for final of 100 mM Tris. After 5 min, the reactions were spun through Zeba 7 kDa columns pre-equilibrated in PBS. The flow through was measured on a nano drop using A280 with IgG setting. Final antibody concentrations were ~ 1.3 mg/mL. One volume of glycerol was added to achieve ~ 0.65 mg/mL antibody in 50% glycerol. Sodium azide was added to final 5 mM. Fluorophore-conjugated antibodies were stored in the dark at −20C.

### Biotinylation of upifitamab

Biotinylation was performed analogously to fluorophore conjugation with several modifications. Upifitamab (100 µL, 2 mg/mL) was desalted in a Zeba 7 kDa column equilibrated in PBS. EZ link sulfo-biotin-LC-LC-NHS (1 mg) was resuspended in 150 µL of PBS to achieve 10 mM solution, with 1.6 µL immediately added to 120 µL of desalted upifitamab in PBS. After 30 min, 13 µL of 1 M Tris pH 8.0 was added. After 5 min at room temp, upifitamab was desalted again in Zeba 7 kDa column pre-equilibrated in PBS. Final upifitamab-biotin concentration was 1.4 mg/mL as measured by nanodrop at 280 nm. Antibody was diluted with one volume of glycerol and sodium azide added to final 5 mM. Biotinylated upifitamab was stored at −20C.

### Flow cytometry of HEK293T cells

HEK293T cells were isolated in versene solution at 37C for 15 min to facilitate cell dissociation to individual cells. Cells were washed and resuspended in PBS + 2 mM EDTA at 1 × 10^7 cells/mL. Cells were incubated with live-dead NIR viability dye (1:2000) at room temp in the dark for 15 min with intermittent gentle vortexing. One volume of FACS buffer (PBS, 2 mM EDTA, 1% BSA) was added and cells centrifuged at 500g x 3 min. Staining was then performed with Alexa647-conjugated Isotype control IgGk or upifitamab on ice for 1h. Cells were washed x2 with FACS buffer, filtered through a 30 µm FACS filter tube and analyzed on a Sony SH800 flow cytometer.

### Mouse breeding and handling

Male and female mice between 6-24 weeks were used for experiments. All related protocols were approved by the Duke University Institutional Animal Care and Use Committee in accordance with NIH guidelines. The following transgenic mice were utilized: *Sftpc-Gfp* (constitutive GFP-expressing under AT2 promoter)^31,32^, *Sftpc tm1(cre/ERT2)Blh (Sftpc-CreER)* (stock number 028054, Jackson Laboratory) and *Rosa26R-CAG-lsl-tdTomato (Ai14)* (stock number 007914, Jackson Laboratory) mice were maintained on a C57BL/6J background. To induce TdT expression in AT2 cells, 3 doses of tamoxifen (0.2 mg/g body weight; 20 mg/mL in corn oil, Sigma Aldrich) were administered intraperitoneally on 3 consecutive days (100 µL per 20 g body weight). Mice were harvested 2 weeks after the final dose.

### Mouse lung harvesting and dissociation for flow cytometry or MACS

Following CO2 euthanasia, the inferior vena cava was incised and 5 mL of PBS perfused via the right ventricle. Lung lobes were removed, washed in PBS, chopped into tiny pieces and cryopreserved in FBS with 10% DMSO (v/v) or immediately dissociated. For each adult mouse, dissociation was performed in HBSS with Ca2+ and Mg2+ containing collagenase I (0.2 mg/mL), Dispase (5 Caseinolytic U/mL) and Zymo recombinant DNAse I (5 U/mL) at 37C with continuous rotation. After 10 min, pieces were pipetted up and down through a P1000 then digested for another 10 min at 37C. One volume of DMEM + 10% fetal bovine serum was added. Dissociated cells were passed through a 70 µm filter, diluted in 2 volumes of FACS buffer and collected by centrifugation. The cell pellet was resuspended in 1 mL of ACK lysis buffer at room temp for 2 min, 4 mL of DMEM added and cells counted. Cells were pelleted by centrifugation at 500 g and resuspended as indicated for flow cytometry or MACS.

### Mouse lung harvesting and processing for IF

Following CO2 euthanasia and perfusion with PBS, lungs were inflated with 1 – 1.5 mL of 4% PFA via the trachea and suture ligated. The tracheobronchial tree with lungs was removed, washed in PBS and fixed in 4% PFA for 3h at RT. Lobes were removed, transferred to 30% sucrose overnight at 4 C, equilibrated in room temp OCT then frozen in OCT blocks. Lungs were cryosectioned at 6 – 8 µm and stored at −40 C.

### Human lung harvesting and dissociation for FACS

Healthy human lung tissue was obtained in accordance with the Duke University Institutional Review Board. Pleura was trimmed away from peripheral lung pieces, which were then chopped and either immediately processed or cryopreserved in fetal bovine serum containing 10% DMSO (v/v). Human lung was dissociated identically to mouse lung, with the exception of higher DNAse 1 concentration (25 U/mL).

### Upifitamab based isolation of AT2 cells by FACS

Mouse and human lung samples were processed identically with the exception of antibodies used for mouse vs human reactivity (e.g., CD45, EpCAM). All centrifugation steps were performed at 500g x 3 min. Following dissociation and RBC lysis, cells were counted and processed at density of 1×10^7^ cells/mL. For flow analysis (not FACS), viability staining was performed identically to HEK293T cells using either zombie violet or live/dead NIR dye. Antibody staining was performed in FACS buffer on ice for 45 min with gentle vortexing every 10 min. Upifitamab was used at concentration of 2 µg/mL unless otherwise indicated. Following live cell staining, cells were strained through a 30 µm FACS tube filter. For cell sorting, viability stain (Helix NP Blue) was added after antibody staining (in lieu of fixable dyes like zombie violet). Flow cytometry was performed on a Sony SH800 analyzer-sorter or Cytek Aurora 5 laser analyzer. Following established scatter and singlet gates, EpCAM+/NaPi2b+ or CD45-/NaPi2b+ cell populations were sorted into DMEM containing 10% FBS, Anti-Anti and 10 µM Y-27632. Sorted cells were centrifuged at 500g for 5 min then immediately counted and seeded into 3D Matrigel domes as described. Flow cytometry data were exported in fcs format and analyzed in FlowJo11.

### Upifitamab based isolation of AT2 cells by MACS

For MACS, cells were processed at density of 0.5-1×10^8^ cells/mL in MACS buffer (identical to FACS buffer except de-gassed prior to use). All centrifugation steps were performed at 500g x 3 min. For 3×10^7^ cells (from two adult mice), cells were suspended in 270 µL of MACS buffer and 30 µL FcR block added. After 5 min, upifitamab-biotin (4.2 µg) was added to achieve a target concentration of 0.15-0.25 µg upifitamab per 10^6^ cells. After rotation at 4 C x 20 min, cells were washed twice with 1 mL MACS buffer then resuspended in 270 µL MACS buffer with 30 µL anti-biotin magnetic beads. Cells were rotated at 4 C x 20 min, washed x3 with MACS buffer, filtered through a 30 µm Celltrix filter into a 1.5 mL microfuge tube, immediately passed through a Miltenyi MS column, washed with 500 µL MACS buffer x3, removed from the magnet and eluted in 500 µL MACS buffer. Yields were generally 0.8-1.2 × 10^6^ cells per adult mouse.

### Culture of mouse and human AT2 cells

Culture of AT2 cells was performed as described using serum-free and feeder cell-free culture conditions^33^. In brief, 2000-5000 MACS-isolated AT2 cells or 5000-10000 FACS-isolated AT2 cells were seeded per 50 µL dome containing 50% Matrigel. Gentamicin (50 µg/mL) was include for the first 3 days of culture. Media was changed every 3 days. Mouse AT2 cells were kept in culture for 10-12 days while human AT2 cells were cultured for 16-20 days between passages. For 2D culture, mouse AT2 cells were seeded to coverslips pre-coated with collagen I (to maintain AT2 identity) or thin-coated with fibronectin to induce AT1 differentiation over 7-10 days as previously described^34^. Cells were allowed to expand for 7-10 days then washed with PBS x1, fixed in 4% PFA for 10 min at room temp then subjected to immunostaining.

### Processing of mouse organoids for microscopy

For imaging, Matrigel domes were cast onto Mat-Tek 35 mm tissue culture plates containing 1.5 mm coverslips. After 10 days in culture, domes were washed with PBS x2, fixed in 4% PFA at 37 C x 30 min, permeabilized with PBS + 0.2% Triton X-100 for 30 min at room temp, then blocked and immunostained overnight. Secondary antibodies were applied for 3 h along with 300 nM DAPI followed by washed x3 with PBS containing 0.1% Tween-20 (PBST). To preserve organoid structure, confocal imaging was immediately performed on the 35 mm dish. Alternatively, the cover slip was removed and mounted to a slide although this causes deformation (flattening) of the organoids.

### Processing of human organoids for microscopy

Human organoids grown in 3D culture were recovered from Matrigel by pipetting up and down with a wide bore P1000 pipette tip, fixed in 4% PFA for 20 min at RT, permeabilized with PBS + 0.1% Triton X-100 for 20 min at RT and subjected to immunostaining in PBST + 1% BSA (w/v) with overnight incubation for primary antibodies. Secondary Ab staining was performed for 2h at RT. During the 3^rd^ and final wash step, 300 nM DAPI was added for 10 min at RT to stain nuclei.

### Confocal microscopy

All imaging was performed on an Olympus FV4000 confocal microscope using Galvano scanning at 2048 × 2048 resolution. Images were processed in Olympus Fluoview and exported as 24-bit RGB in TIFF format.

## RESULTS

### Surfaceome analysis supports NaPi2b as a candidate AT2 surface protein

It was hypothesized that published transcriptomic datasets could be leveraged to identify AT2 surface proteins amenable for live cell enrichment. To obtain a candidate list of proteins, human and mouse proteomes were filtered to include only proteins with potential for cell surface expression, specifically those included in the established SURFY in silico database (human “surfaceome”)^20^ of 2886 proteins. A total of 5 mouse and 4 human datasets of healthy lung were included in this surfaceome analysis (Figure S1). As shown in Figures 1A-B, the sodium phosphate transporter NaPi2b (gene product SLC34A2) exhibits a highly AT2-restricted expression pattern across all analyzed datasets. This multi-pass transmembrane protein resides on the apical surface of AT2 cells where it plays a critical role in sodium phosphate transport^35^ from the alveolar lumen into AT2 cells, thus preventing pathologic accumulation of insoluble phosphate salts in the alveolar space^36,37^. Flow cytometry for NaPi2b has been used to label AT2 cells^3^. However, antibodies against NaPi2b generally target the cytoplasmic C-terminus^38,39^ thus requiring cell permeabilization that is incompatible with live-cell isolation methods. By virtue of its AT2-specificity and apical/surface localization, NaPi2b was carried forward as the primary target to improve AT2 cell isolation methods.

**Figure 1.**
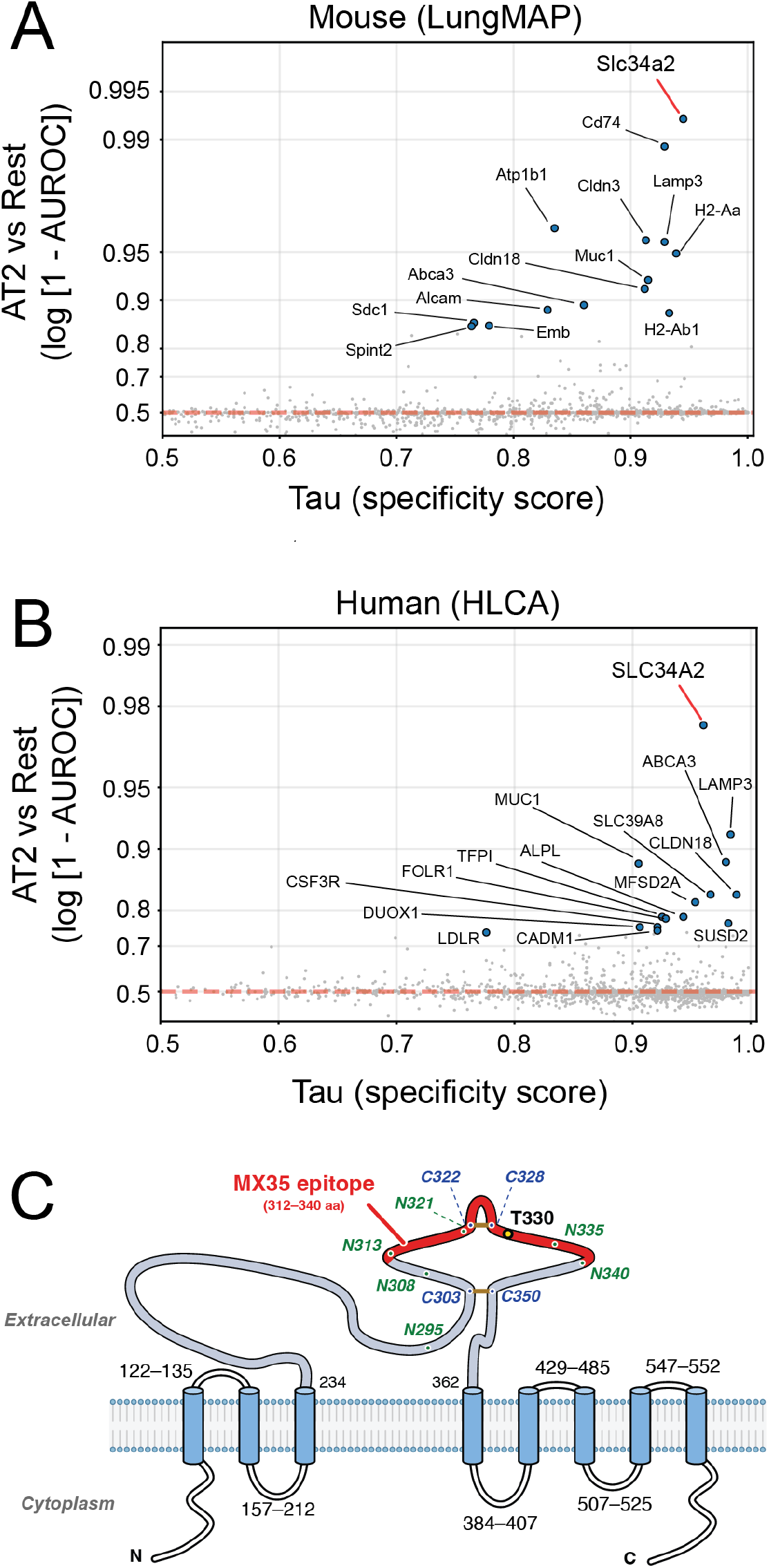
Surfaceome analysis of mouse and human transcriptomes identifies NaPi2b (Slc34a2) as a candidate AT2 surface marker. Scatterplots show tau score on x-axis (indicative of overall transcript specificity for a single cell type with range 0-1) and relative expression of each gene in AT2 cells compared to all other annotated cell types on y-axis. Shown are two representative published datasets from (**A**) LungMap Mouse and (**B**) Human Lung Cell Atlas (HLCA). Each data point is a gene whose protein product is included in the SURFY human cell surface proteome (2886 proteins). Non-surfaceome proteins are omitted from the analysis as they would not support live cell isolation. (**C**) Schematic of sodium phosphate transporter NaPi2b. The antibody-targeted second extracellular domain (ECD) is expanded. Amino acids 312-340 are targeted by mouse monoclonal MX35 and commercial derivatives to varying extents. Green and blue numbers denote sites of potential glycosylation and disulfide bond formation, respectively. The C303-C350 disulfide is predicted by UniProtKB while C322-C328 is predicted by AlphaFold (shown in brown). Residue T330 is critical for MX35 binding as mutation at that residue causes loss of antibody recognition. Residue numbering is specific to human NaPi2b.

### Consideration of upifitamab as a species-agnostic NaPi2b-reactive antibody

Given high level expression of NaPi2b in ovarian and lung malignancies, this protein has received significant attention as a therapeutic target for antibody-drug conjugates (ADCs). Specifically, the second extracellular domain (ECD) was targeted by mouse mAb MX35 to support extensive pre-clinical work on targeting ovarian cancers^40–42^. MX35 was “humanized” to Rebmab200^43^ while analogous industry-led efforts generated Lifastuzumab^44^ and upifitamab^45^. Each of these antibodies targets the 2^nd^ ECD of NaPi2b (Figure 1C).

While each of these antibodies are strongly reactive to human NaPi2b, reactivity to NaPi2b from other species is variable. For example, MX35 is unreactive towards murine NaPi2b at least by western blot^41^, and Lifastuzumab appears unreactive towards rodent NaPi2b due to alteration of human Leu^327^ (Tyr in rodents)^44^. A threonine at position 330 is essential for MX35-derived antibody affinity^46^, although conservation of this residue between human and mouse implies other determinants in species reactivities. In contrast to MX35 and lifastuzumab, upifitamab showed immunoreactivity to synthetic peptides derived from mouse NaPi2b with comparable affinity to human NaPi2b^45^. Thus, it was hypothesized that upifitamab might provide a species-agnostic route for AT2 cell isolation.

### Assessment of upifitamab reactivity towards murine NaPi2b

To determine if upifitamab is reactive to mouse NaPi2b, HEK293T cells were transduced with lentivirus bearing murine NaPi2b with a C-terminal HA tag. Immunofluorescence of fixed cells showed strong reactivity of upifitamab to cells expressing NaPi2b but not to control cells expressing Cyclophilin-A (Figure S2A). Analogous live-cell labeling showed strong surface staining of upifitamab in a dose-dependent and NaPi2b-specific manner by confocal microscopy (Figure S2B) and flow cytometry (Figure S3). Thus, upifitamab indeed binds to murine NaPi2b and is not restricted to the human orthologue like other anti-NaPi2b monoclonal antibodies.

### Assessment of AT2-cell specificity of upifitamab by confocal immunofluorescence

Having established that upifitamab efficiently labels murine NaPi2b ectopically expressed in HEK293T cells, subsequent experiments focused on evaluating the AT2-cell specificity of upifitamab. To this end, C57BL/6 mice bearing AT2 cell-restricted TdTomato (TdT) expression (*Sftpc-CreER / Lox-Stop-Lox TdT*) were treated with tamoxifen to induce TdT in AT2 cells, then harvested and lungs analyzed by confocal immunofluorescence (IF). As shown in Figure 2A, upifitamab strongly labeled the apical surface of TdT+ population (AT2 cells) where NaPi2b is known to localize. Next, upifitamab was compared to EpCAM which is a primary workhorse for AT2 cell isolation. As shown in Figure 2B, upifitamab showed strong overlap with AT2 marker surfactant protein C (SPC) and EpCAM within the alveolar regions. Unlike EpCAM which is highly expressed on airway epithelium, NaPi2b showed restriction to AT2 cells at least under these resting/healthy conditions. Examination of small airways showed a similar pattern where NaPi2b is restricted to AT2 cells and not present in distal secretory/progenitor populations such as club cells and bronchoalveolar stem cells (BASCs) (Figure 2C). In human lung, analogous pattern of NaPi2b expression was characterized using upifitamab, again with strong overlap to SPC-positive cells and restriction to the apical AT2 cell membrane (Figure 2D).

**Figure 2.**
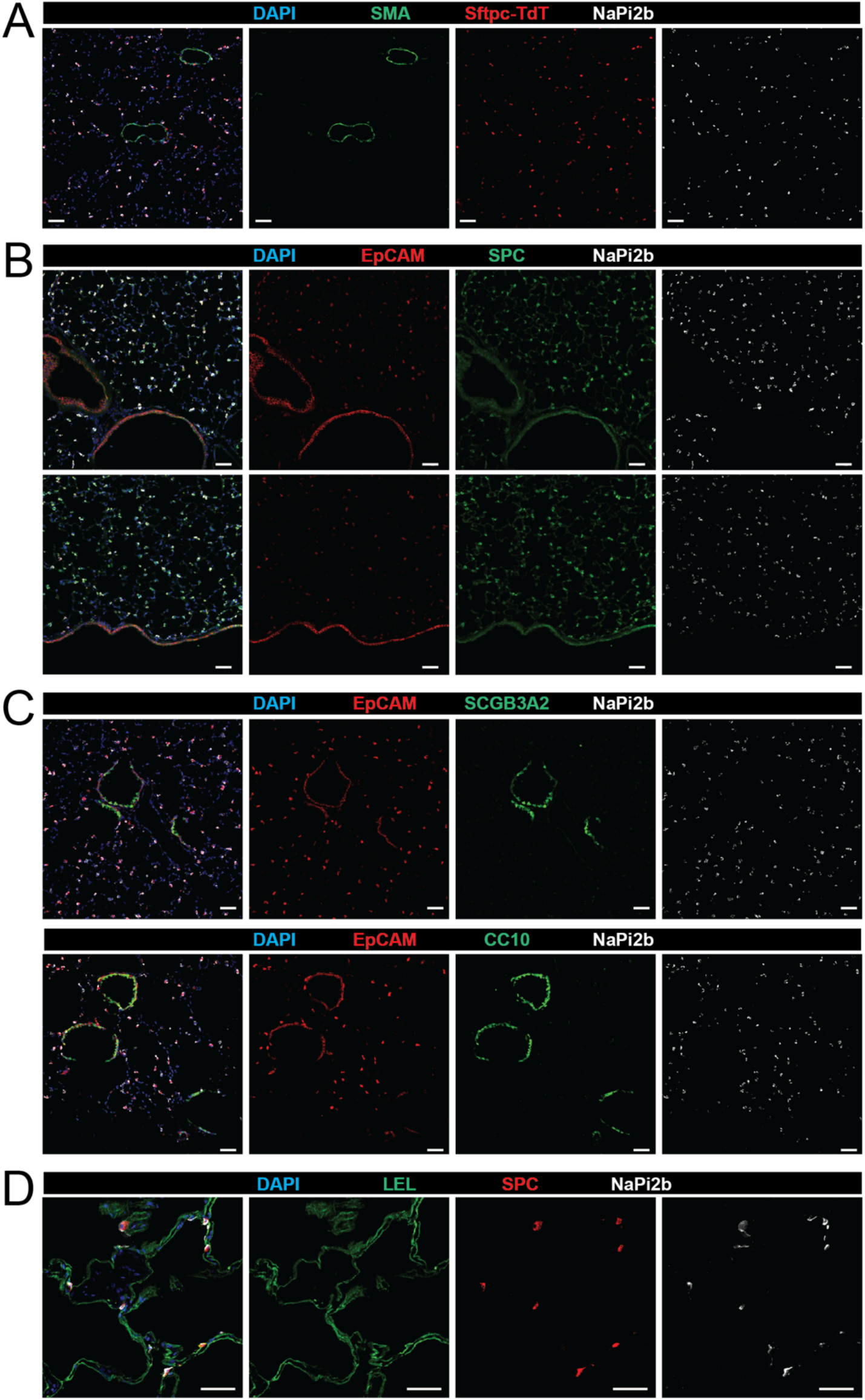
Confocal microscopy of healthy mouse and human lung immunostained with upifitamab. (**A**) Sftpc-CreER/Lsl-TdT mice were treated with tamoxifen to induced lineage labeling of AT2 cells (TdT positive) then harvested, cryosectioned, stained and imaged at 20x magnification. Smooth muscle actin (SMA) indicates peribronchiolar and perivascular smooth muscle. (**B**) Healthy mouse lung co-stained for EpCAM and SPC (surfactant protein C, Sftpc) at 20x magnification. Airway epithelium are the strongly EpCAM+ cells lining the airway lumens. (**C**) Analysis of upifitamab staining specificity in small airways at 20x magnification. Club cells and bronchoalveolar stem cells (BASCs) were immunostained with antibodies against SCGB3A2 and CC10 (SCGB1A1). (**D**) Healthy human lung immunostained and imaged at 40x magnification with max intensity projection (3 slices x 1.4 µm z-steps). Lycopersicon esculentum lectin (LEL) reacts with the alveolar surface of AT1 cells. In each panel, NaPi2b is immunostained with Alexa647-conjugated upifitamab. Data are representative of at least four healthy adult mice and four healthy human lung donors. Scale bar = 50 µm.

### Application of upifitamab for mouse AT2 cell discrimination by flow cytometry

Next, lungs from mice with constitutive AT2-cell specific expression of GFP (Figure S4) were used to assess upifitamab’s efficacy in AT2 labeling for flow cytometry. Following standard gating strategies for single viable cells (Figure 3A), upifitamab showed strong enrichment of GFP+ cells (i.e., AT2 cells) when coupled to CD45-gating (Figure 3B) or EpCAM+ selection (Figure 3C) relative to isotype control IgGk. Similarly, gating first for GFP+ cells (AT2 cell lineage) demonstrated an expected reciprocal pattern of EpCAM and NaPi2b positivity (Figure 3D). As summarized in Figure 3E, these data support at least two gating strategies to obtain highly specific (>99%) AT2 cell labeling, which include CD45-/NaPi2b+ or EpCAM+/NaPi2b+ populations. With regards to recall (i.e., yield of AT2 cells retrieved), the CD45-/NaPi2b+ strategy showed an ~15% advantage over EpCAM+/NaPi2b+ approach. These gating approaches can be combined (i.e., CD45-/EpCAM+/NaPi2b+) while still preserving a simple overall workflow (Figure S5). Alternatively, EpCAM can be replaced by MHCII subunit IA/IE (Figure S6), although this approach relies on negative gating of immune cells and is not compatible with human AT2 cells (mouse restricted)^12^.

**Figure 3.**
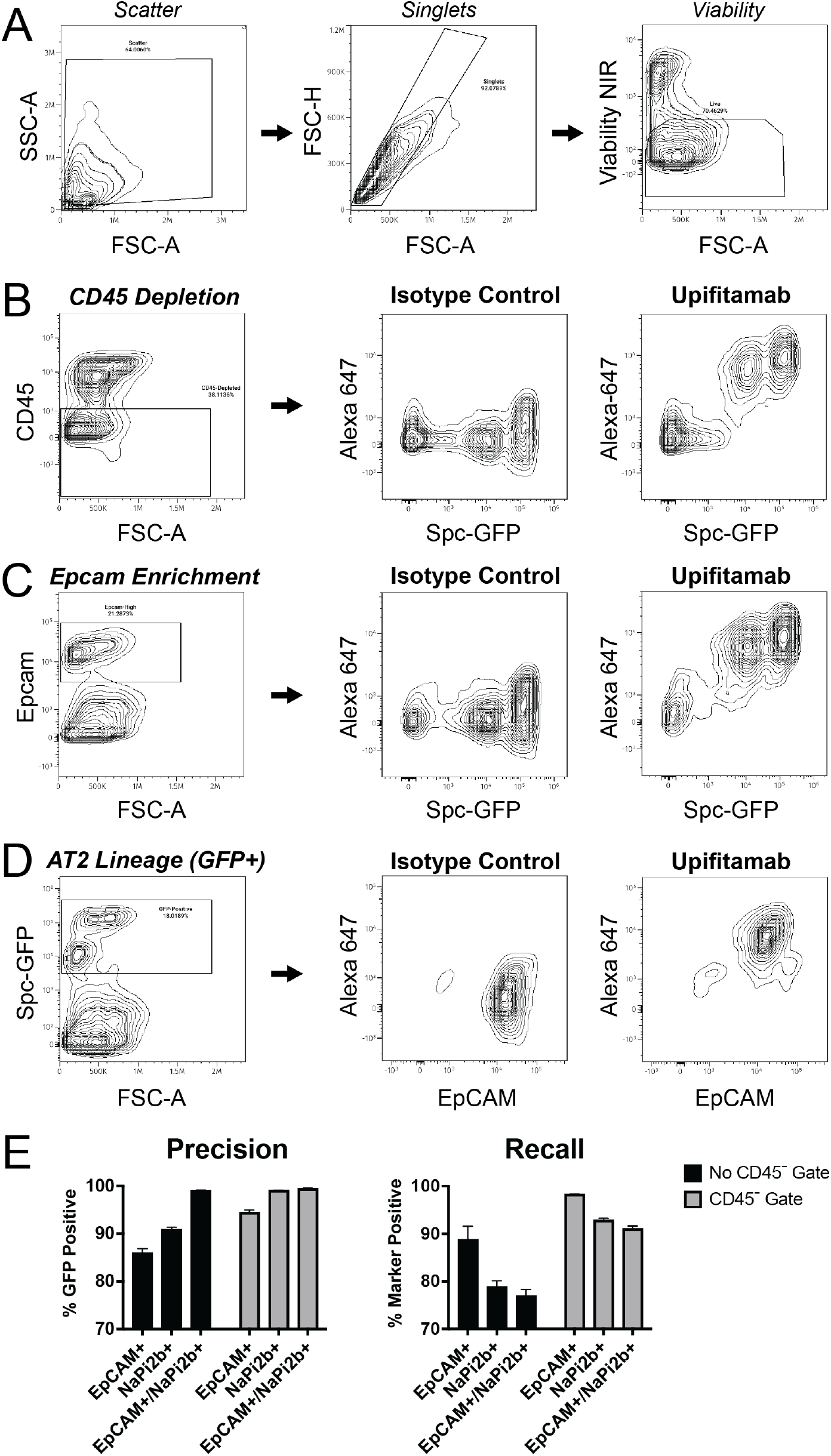
Upifitamab allows for simple AT2 cell identification by flow cytometry. Lungs from *Sftpc-Gfp* mice (harboring AT2-specific GFP expression) were dissociated and subjected to live cell staining followed by flow cytometry. (**A**) General gating strategy including scatter, singlets and viability. (**B**) Gating for CD45 negative population, followed by lineage label (GFP) on x axis and isotype control vs upifitamab on the y-axis. (**C**) Gating for EpCAM positive population, followed by similar plot as in (**B**). These two plots assess specificity of upifitamab for AT2 cells with two different co-gating strategies. (**D**) Gating for AT2 cells (GFP+) followed by plots of EpCAM on the x-axis and isotype control IgGk vs upifitamab on the y-axis. The GFP+ population is overwhelmingly EpCAM and NaPi2b positive. (**E**) Assessment of precision and recall for AT2 isolation by various strategies. For each n = 4 independent experiments.

### Functional assessment of mouse AT2 cells obtained by upifitamab-coupled FACS and MACS

Having established methods employ upifitamab for live cell AT2 cell labeling, healthy mouse AT2 cells were isolated by both FACS and MACS followed by assessment by cell culture. By FACS, typical recoveries were between 0.3 – 0.5×10^6^ cells per adult mouse. By MACS, yields were significantly higher between 0.8 – 1.2×10^6^ cells per adult mouse. Following established culture methods^33^, AT2 cells were seeded to domes in 50% Matrigel and allowed to expand in 3D organoid culture. Organoids showed expected morphologies and general growth rates (Figure 4A). Compared to FACS, MACS-isolated mouse AT2 cells expanded earlier in the culture period, exhibiting spheres as early as days 4-5. After 10 days in culture, domes were fixed in PFA, permeabilized, immunostained and analyzed by confocal imaging. Markers of AT2 cells showed strong and global expression within intact whole organoids (Figure 4B) and thinner organoid fragments (Figure 4C), thus confirming proper cell identity. Following 3D expansion, AT2 cells were seeded to 2D culture on a thick coat of collagen I or thin coat of fibronectin to preserve AT2 cell identity or to induce differentiation to AT1 cells, respectively^34^. As expected, cells grown on collagen I maintained AT2 cell morphology and marker expression (Figure 4D) while those grown on fibronectin-coated glass differentiated to flat AT1 cells within 1 week of culture (Figure 4E). These results establish that upifitamab allows for rapid and simple AT2 cell isolation from mice, thus supporting AT2 cell culture experiments. While MACS provided higher AT2 cell yields, overall identical cell culture results were obtained by the two methods.

**Figure 4.**
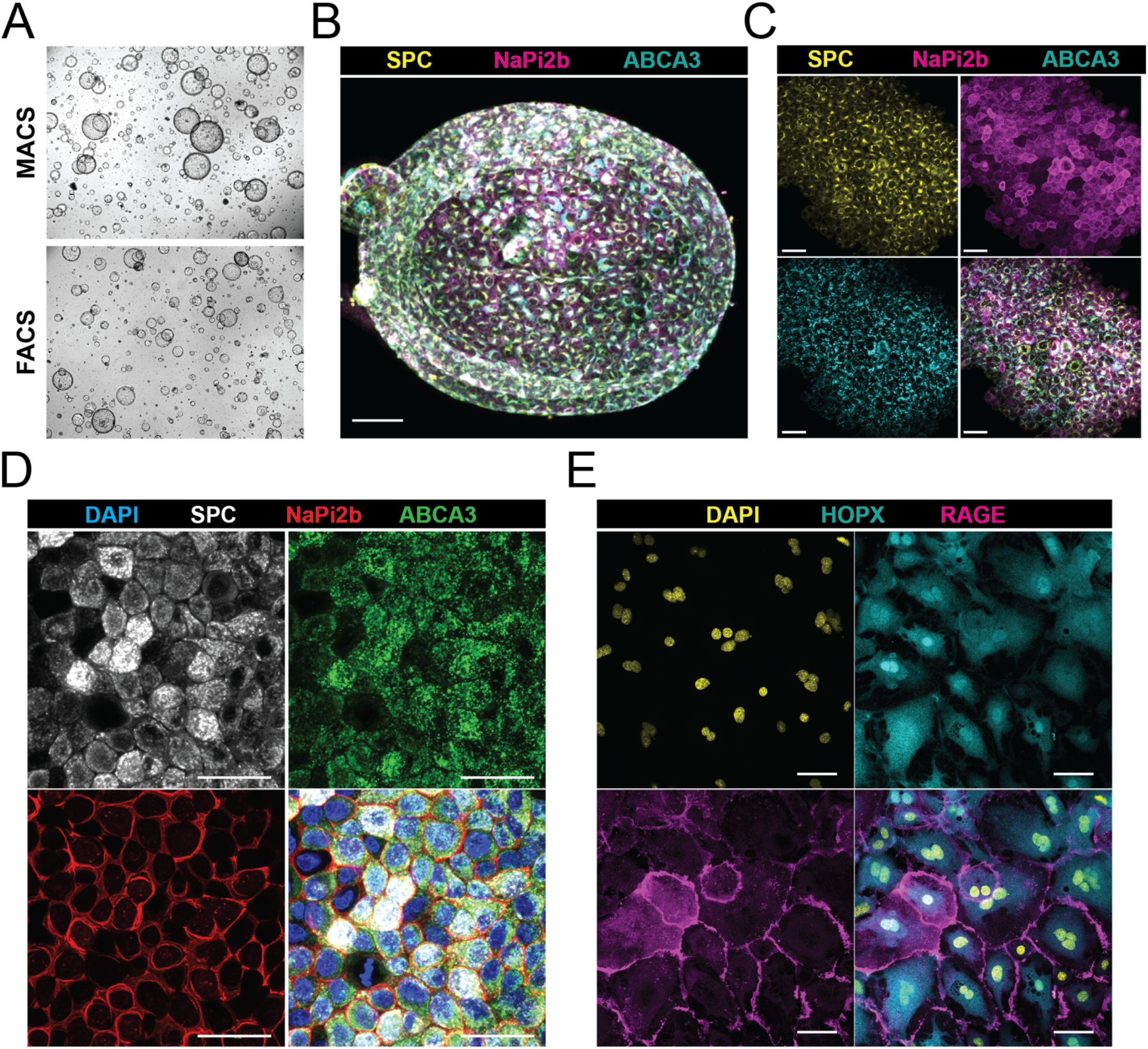
Culture of murine AT2 cells isolated by upifitamab-coupled FACS and MACS. (**A**) AT2 cells were isolated from healthy mouse lung by FACS (EpCAM+/NaPi2b+ population) or MACS (NaPi2b+ selection), expanded in Matrigel domes via 3D organoid culture then imaged on day 7 by brightfield microscopy at 4x magnification. (**B**) Confocal microscopy of intact day 10 organoid at 20x magnification with maximum intensity projection (MIP) across 40 slices using 1.48 µm z-steps. Scale bar = 100 µm. (**C**) Confocal microscopy of day 10 organoid at 40x magnification with MIP across 15 slices using 1.0 µm z-steps. Scale bar = 50 µm. (**D**) AT2 cells seeded to 2D culture on thick-coat collagen, grown for 7 days then imaged by confocal microscopy at 60x magnification. Four slices using 1.0 µm z-steps were used for MIP. Scale bar = 40 µm. (**E**) AT2 cells were seeded onto a thin coat of fibronectin and allowed to differentiate to AT1 cells over 7 days, then imaged by confocal microscopy at 40x magnification. Scale bar = 50 µm.

### Upifitamab allows isolation and culture of human AT2 cells

To assess whether similar strategies could be used to isolate human AT2 cells, healthy human lung was subjected to live cell staining and flow cytometry. As shown in Figure 5A, viable single cells were positively gated for CD31-/CD45-population then further analyzed for AT2 markers. A large subgroup of the EpCAM+ population showed strong co-expression of NaPi2b as assessed by upifitamab binding, which was not observed with isotype control IgGk. This group represents ~80% of EpCAM+ cells, which correspond to AT2 cells while avoiding other EpCAM+ non-AT2 cells such as airway epithelium. Next, a simple FACS strategy requiring negative gating of CD45 and positive gating of NaPi2b was used to flow sort AT2 cells (Figure 5B). These cells were seeded in 50% Matrigel domes and expanded in 3D organoid culture, then imaged by brightfield microscopy (Figure 5C). Compared to mouse organoids which are spherical, the human organoids exhibited higher cell density with a grape-like morphology. Human organoids were recovered from Matrigel, PFA-fixed, immunostained for established AT2 markers and imaged by confocal IF (Figure 5D).

**Figure 5.**
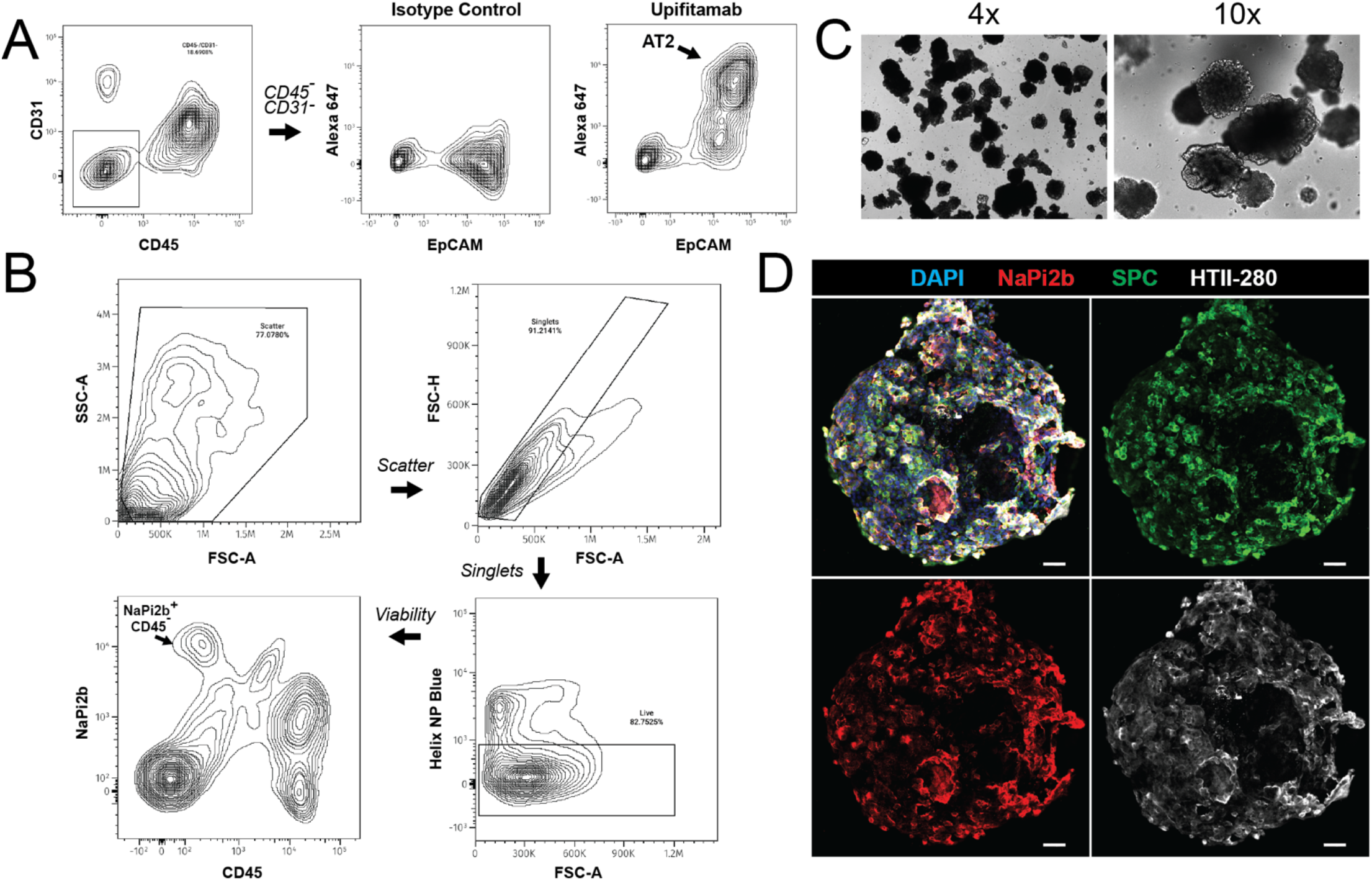
Upifitamab allows for isolation of human AT2 (hAT2) cells. (**A**) Comparison of Alexa647-conjugated isotype control and upifitamab by flow cytometry on healthy human lung. (**B**) Simplified gating strategy for AT2 cell isolation from healthy human lung requiring only gating out CD45+ cells and positive selection for NaPi2b. (**C**) Culture of hAT2 cells in 3D Matrigel at day 17 imaged by brightfield microscopy at 4x and 10x magnification. (**D**) Confocal IF of human organoid stained with apical-specific Ab HTII-280, SPC and NaPi2b. Shown is an MIP of 3 slices at 1.0 µm z-step. Scale bar = 50 µm. For 3D culture, 2500 cells were seeded per Matrigel dome. Results are representative of three separate experiments.

## DISCUSSION

Alveolar type 2 cells are critical for many aspects of alveolar homeostasis and their dysfunction is highly linked to lung disease. Accordingly, AT2 enrichment methods are central to *in vivo* and *ex vivo* studies of lung biology, as well as AT2 isolation for future therapeutic purposes. While methods are established for AT2 cell isolation, to date none are globally applicable across species while requiring distinct antibody combinations or reagents for mouse AT2 vs human AT2 isolation. For example, most mouse AT2 cell isolation strategies rely primarily on gating of EpCAM+ cells in conjunction with other markers to enhance AT2 specificity, whereas human AT2 cell isolations utilize completely different approaches such as surface labeling of HTII-280. Further, chemical reagents such as lysotracker are not amenable to MACS, while other markers such as EpCAM or IA/IE also label other cell types such as airway epithelium and immune cells, respectively. In contrast, the upifitamab-based approach is simple, versatile, species-agnostic and readily adaptable between isolation technologies (i.e. MACS and FACS compatible). In our hands, upifitamab-based FACS yields robust AT2 cell numbers (~300-400k per adult mouse) while MACS provides at least twice the number of AT2 cells (>800k per adult mouse). From a supply standpoint, upifitamab was produced in large quantities (for clinical trials), is commercially available and can be recombinantly expressed.

Despite the utility of this approach, the use of upifitamab for AT2 cell isolation raises some critical questions. First and foremost, does NaPi2b expression change in conditions of injury or disease? For example, in IPF and other ILDs, canonical AT2 cells decrease with concomitant increases in aberrant transitional and basaloid cell states^47,48^. Under such disease conditions where epithelial boundaries are blurred, established AT2 isolation methods unsurprisingly show diminished AT2 cell purity and increased heterogeneity^14,49^. While this can be variably interpreted as “AT2 marker failure” or “inherent pathologic heterogeneity”, the extent to which such limitations apply to NaPi2b-upifitamab remains to be established. Thus, assessment of disease-specific NaPi2b expression is warranted before upifitamab is applied to AT2 cell isolation in the context of disease states.

Secondly, a minor limitation of using humanized monoclonal Abs in human tissues is the inability to use secondary anti-human Abs (due to endogenous human IgG interference). However, this issue is readily circumvented by fluorophore-conjugating upifitamab.

Notwithstanding these limitations and unanswered questions, this new application of upifitamab overcomes multiple current bottlenecks in AT2 cell isolation. By simplifying and streamlining the isolation of AT2 cells, this methodology should make AT2 cell culture more broadly accessible while accelerating research on alveolar homeostasis and disease.

## RESOURCE AVAILABILITY

For additional details or requests for resources/reagents, please contact the corresponding author. All unique reagents generated in this study are available from the lead contact with a completed Materials Transfer Agreement, including fluorophore-conjugated or biotinylated upifitamab.

## DATA AND CODE AVAILABILITY

All code used for data analyses is available at https://github.com/mt-forrester/SLC34A2_NaPi2b-biorxiv/.

## ACKNOWLEDGMENTS

This work was funded by the National Institutes of Health grants K08HL181195 (to MTF), K08CA263300 (to CEE), R01HL153375 and R01HL160939 (to PRT), R01ES034350 (to RMT), Duke Department of Medicine Chairs Research Award (to MTF), Office of Physician Scientist Development Technician Support Award (to MTF) and Nanaline H. Duke Fund Strong Start Award (to MTF).

## AUTHOR CONTRIBUTIONS

MTF conceived the project, designed and executed experiments, analyzed data and wrote the manuscript. SL, RLF, and KE performed AT2 cell culture and analyzed results; KR harvested and processed mouse lungs; RIC and BL provided guidance and training on flow cytometry; KT, JMM, TJW, AT, CEE, RMT and PRT made intellectual contributions, assisted with experimental design and analyzed data.

## SUPPORTING INFORMATION

### KEY RESOURCES

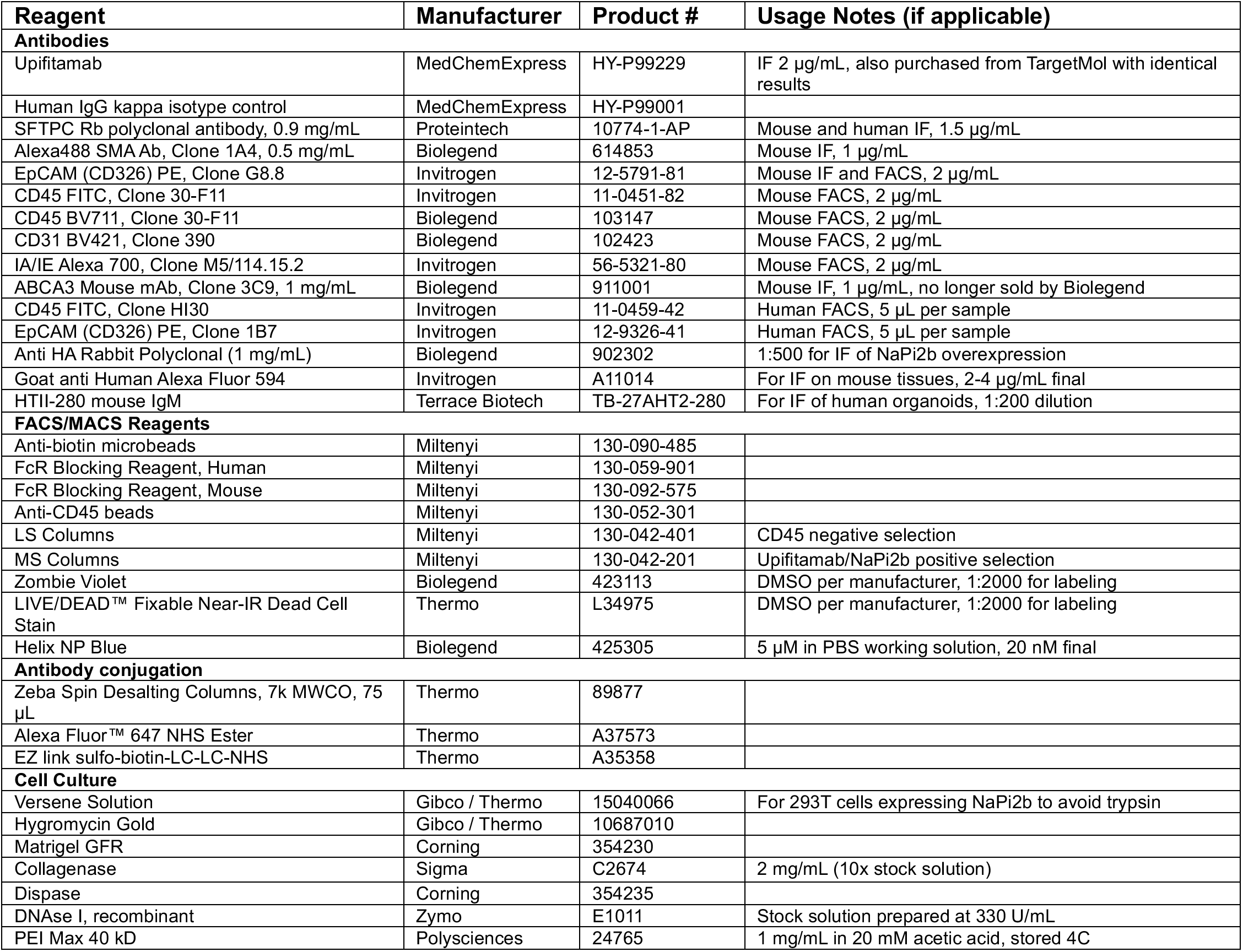

**Figure S1.**
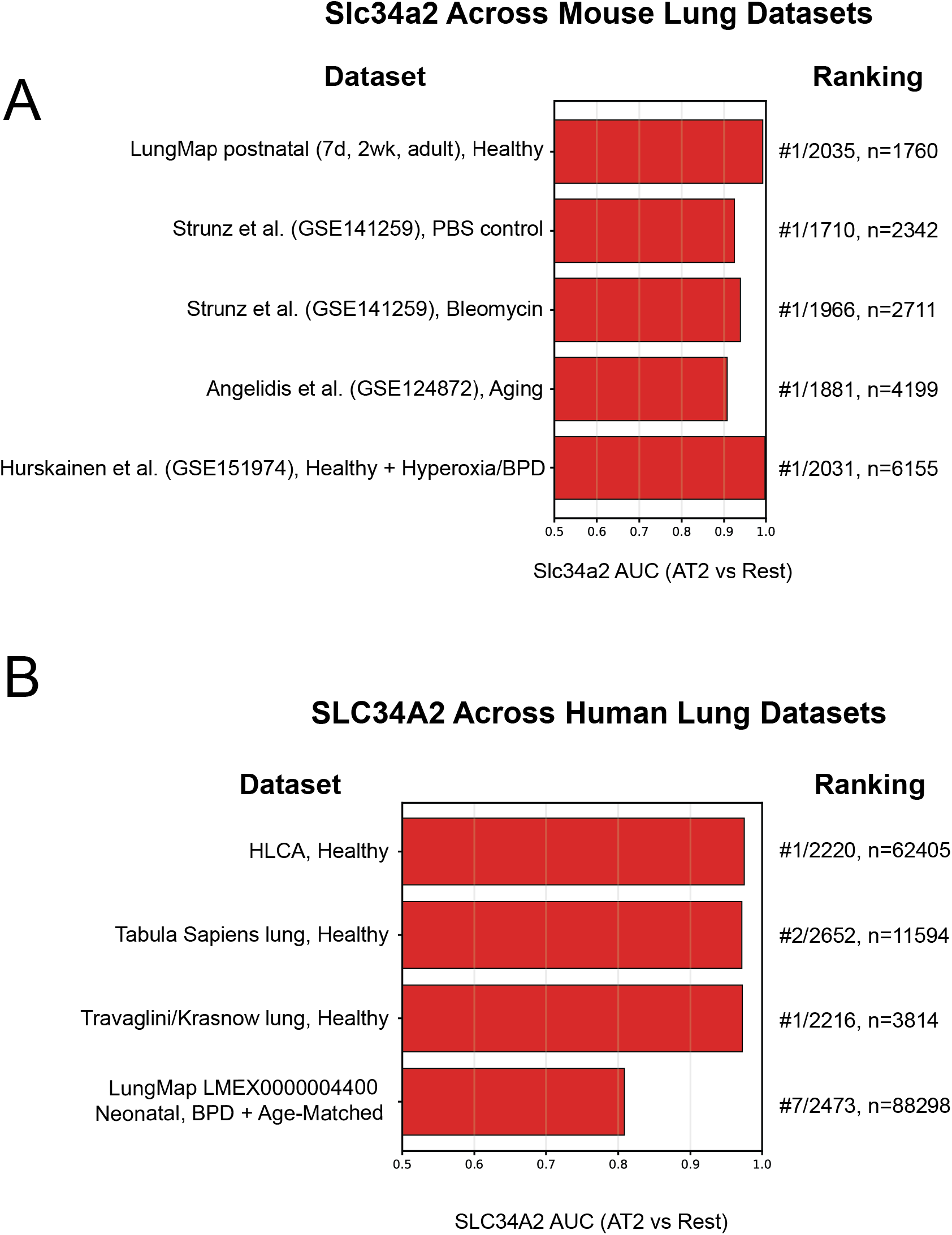
Surfaceome analysis of NaPi2b (SLC34A2) across multiple published transcriptomic datasets. Barplots showing AUROC for NaPi2b transcript from **(A)** Mouse (Slc34a2) and **(B)** Human (SLC34A2). Corresponding dataset is listed to the left of the y-axis. To the right is overall ranking of NaPi2b compared to other surfaceome proteins represented in each dataset followed by the number of AT2 cells included in analysis.

**Figure S2.**
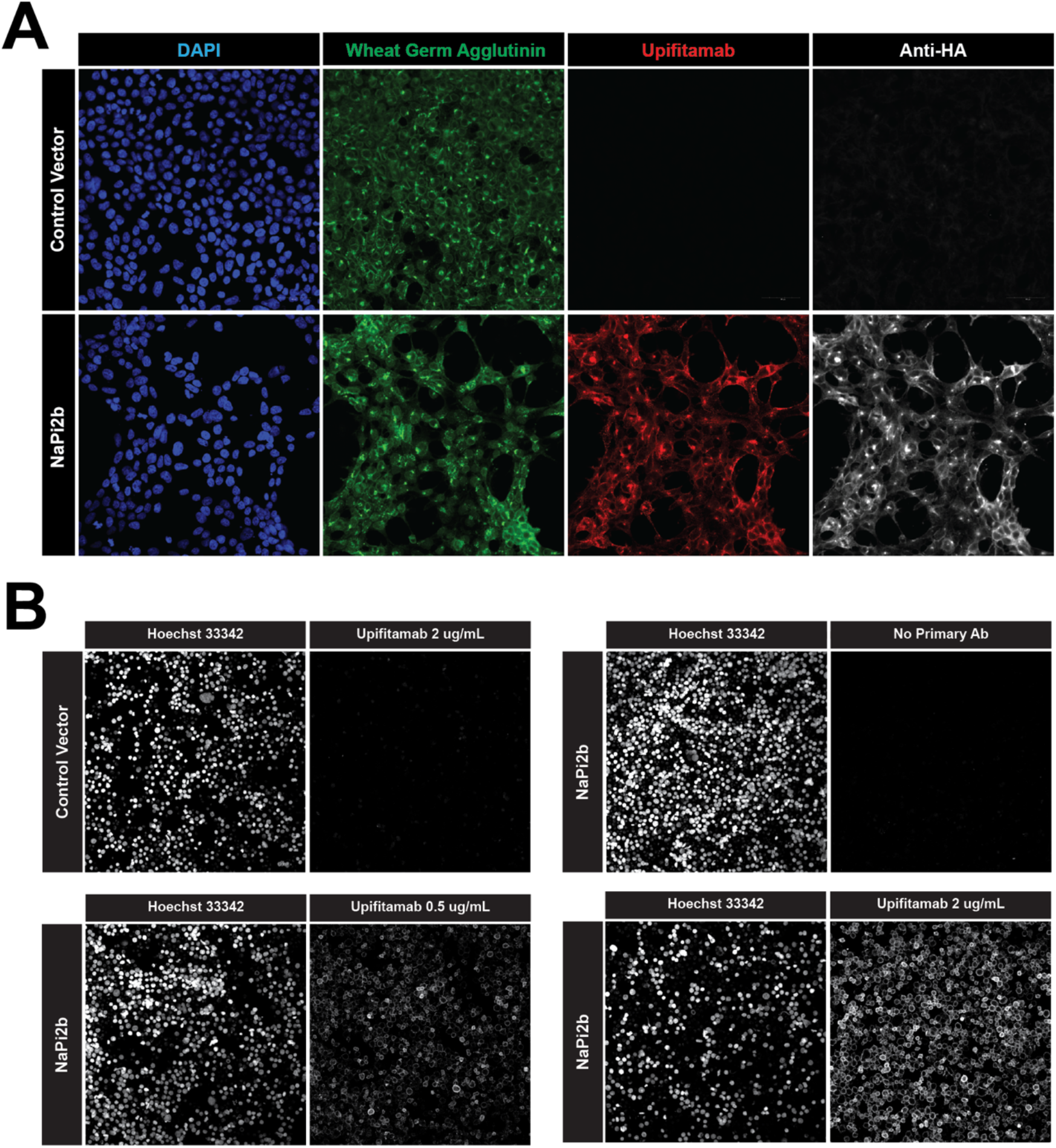
Upifitamab is immunoreactive towards mouse NaPi2b. HEK293T cells were stably transduced with lentiviruses expressing mouse NaPi2b with a C-terminal HA tag or Cyclophilin A (CyA) as a negative control. (**A**) Cells were fixed with PFA, permeabilized and subjected to immunostaining with Upifitamab and rabbit anti-HA antibody to serve as a positive reference signal. Membranes were stained with wheat germ agglutinin (WGA) conjugated to Alexa Fluor 488. (**B**) HEK293T cells were collected with versene (to avoid trypsinization of surface proteins) and subjected to live-cell staining with Upifitamab. Cells were imaged by confocal microscopy at 20x magnification. Results are representative of 3 separate experiments. Horizontal banners indicate antibody or visualization method. Vertical banners indicate control (Cyclophilin A) vs NaPi2b-HA expressing HEK293T cells.

**Figure S3.**
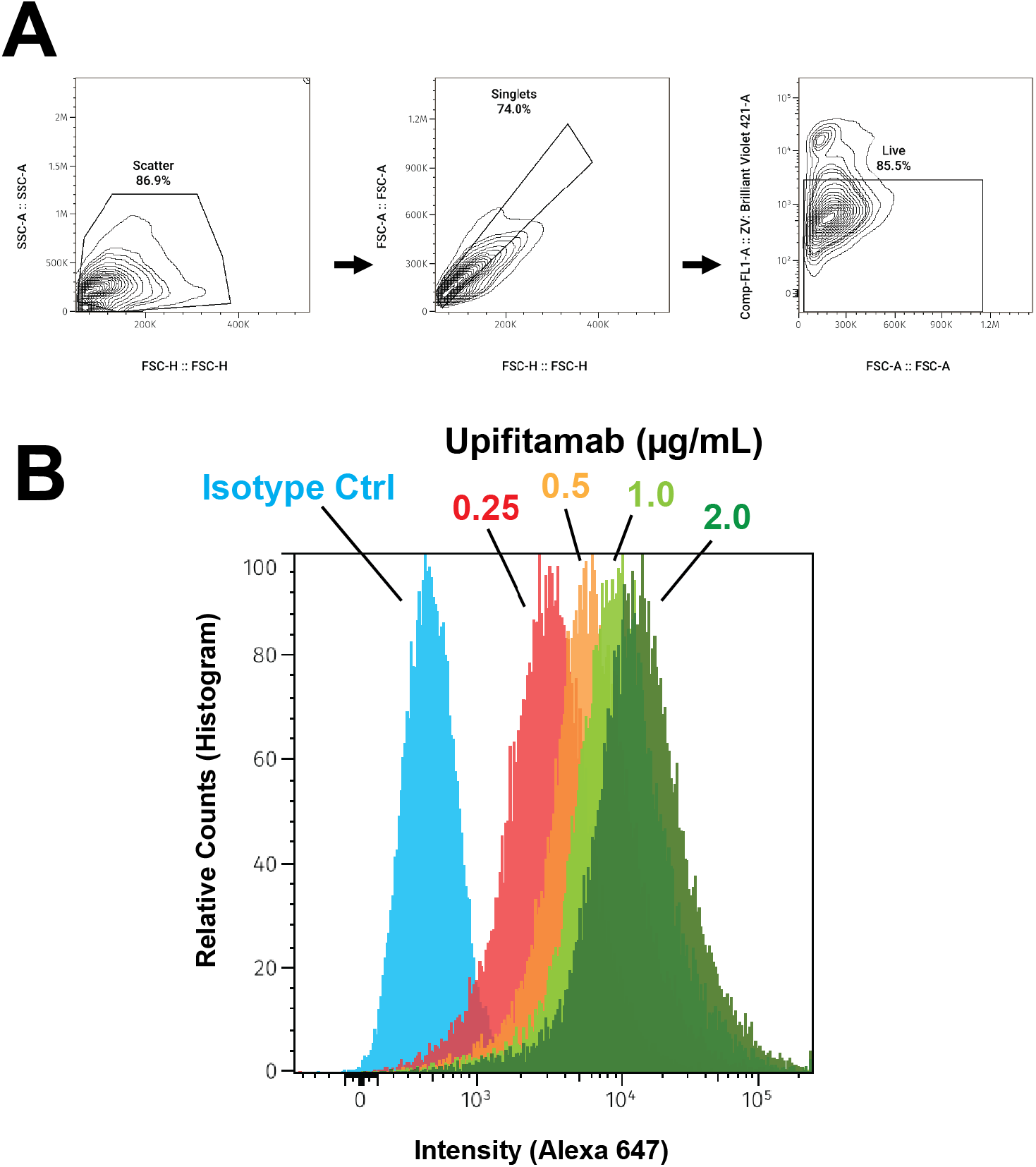
Flow cytometry of stably transduced HEK293T cells establishes specificity of Upifitamab for murine NaPi2b. HEK293T cells were stably transduced with lentivirus expressing either Cyclophilin-A (CyA) or murine NaPi2b (Slc34a2). Following selection with hygromycin, HEK293T cells were collected with versene, processed and analyzed by flow cytometry. (**A**) Gating strategy by scatter, singlets then viability assessed by zombie violet staining. (**B**) Overlaying histograms for Alexa647 intensity comparing isotype control and increasing doses of Upifitamab. Results are representative of four separate experiments. Comparison includes Alexa Fluor 647-conjugated human isotype control IgG kappa vs Upifitamab.

**Figure S4.**
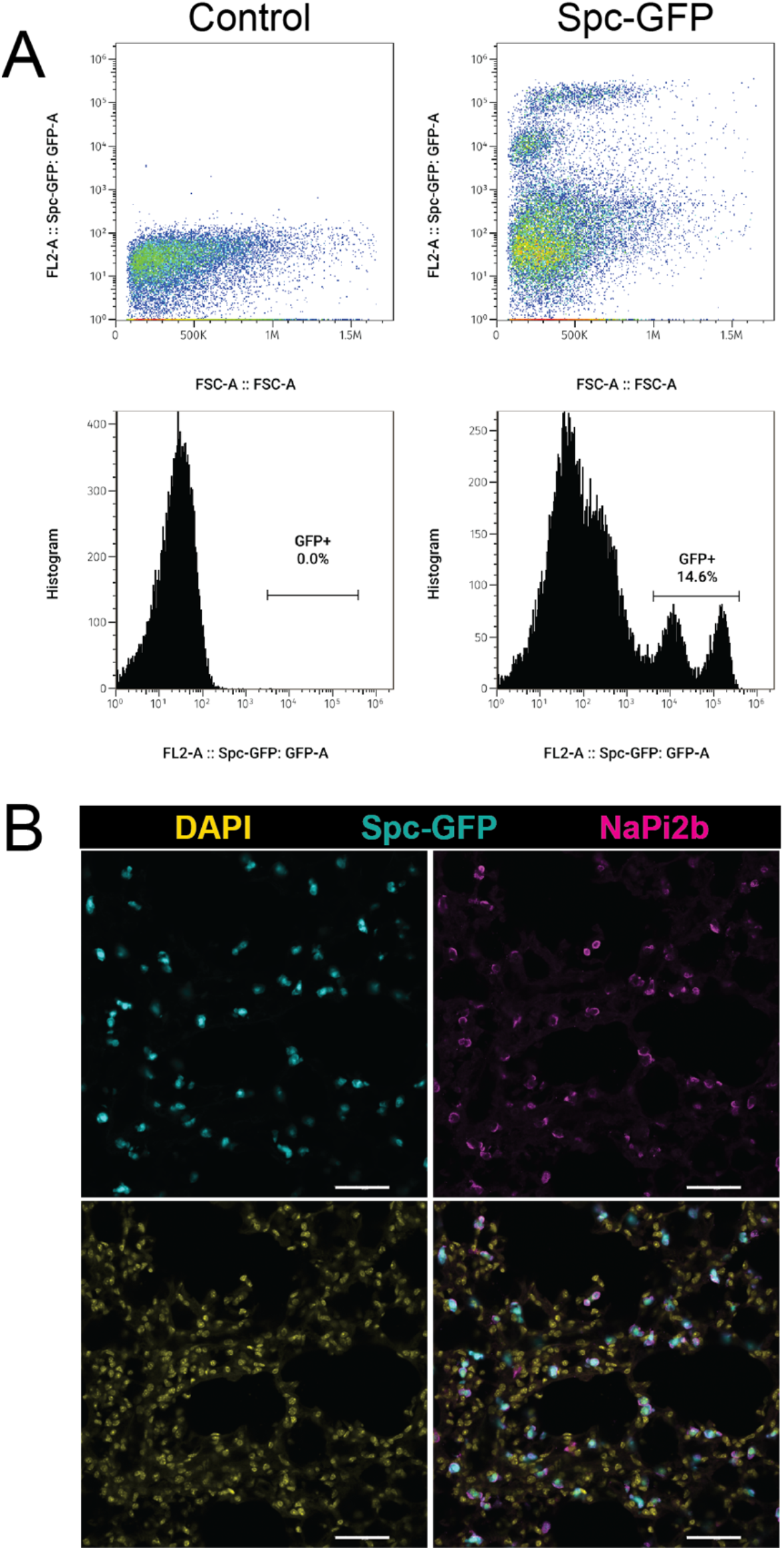
Assessment of Sftpc-GFP (Spc-GFP) mice by flow cytometry and immunostaining with Upifitamab. (**A**) Flow cytometry of compensation controls of whole dissociated cells from unstained WT and Spc-GFP lungs. (**B**) IF and confocal microscopy of Spc-GFP lung at 40x magnification. NaPi2b was stained with Upifitamab. Scale bar = 50 µm. Results are representative of at least 3 separate experiments.

**Figure S5.**
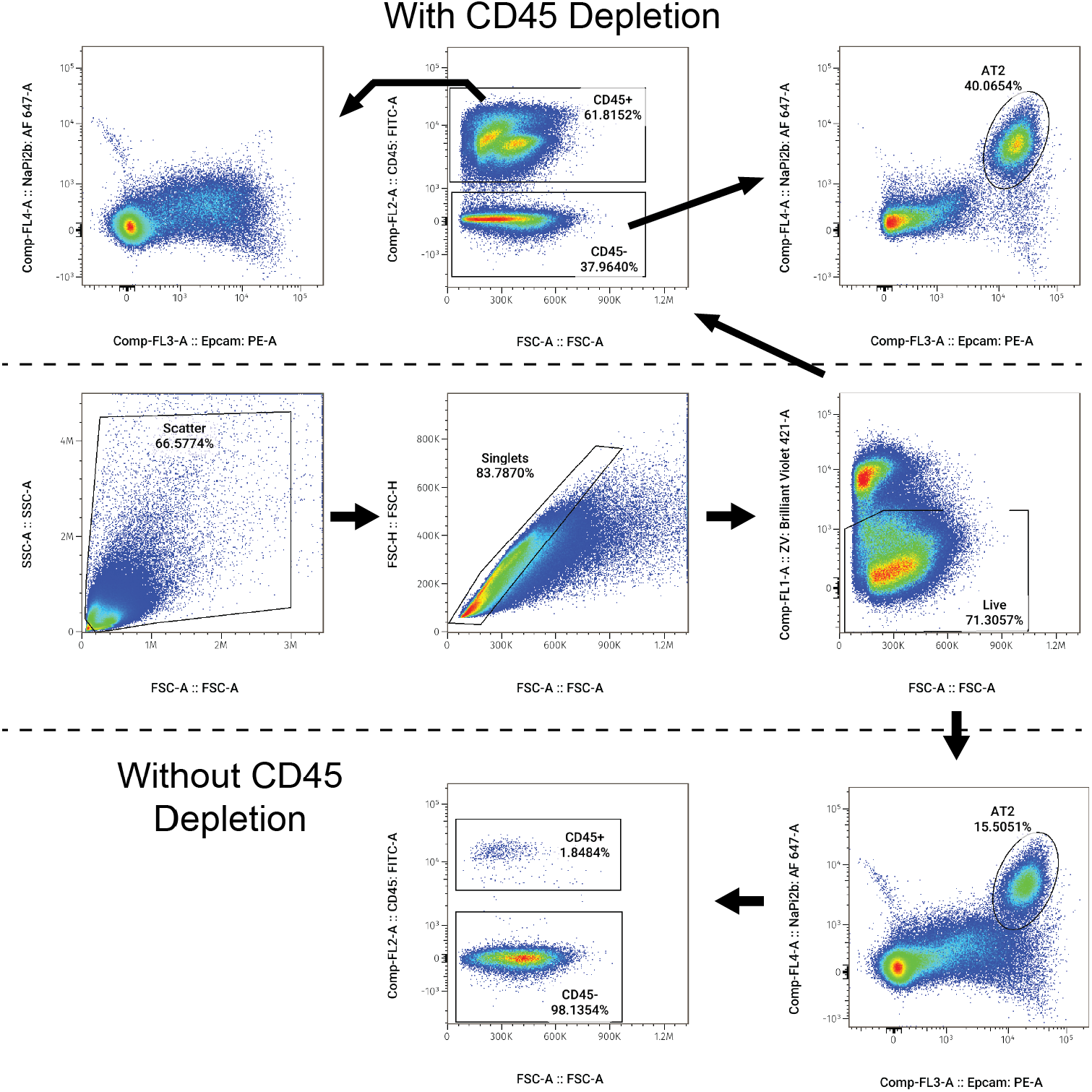
Gating out CD45+ cells provides a modest improvement in flow cytometric resolution of AT2 cells. Mouse lungs were dissociated and subjected to live cell-immunostaining. Data were acquired on a Sony SH800 flow cytometer. Results are representative of n=4 separate experiments.

**Figure S6.**
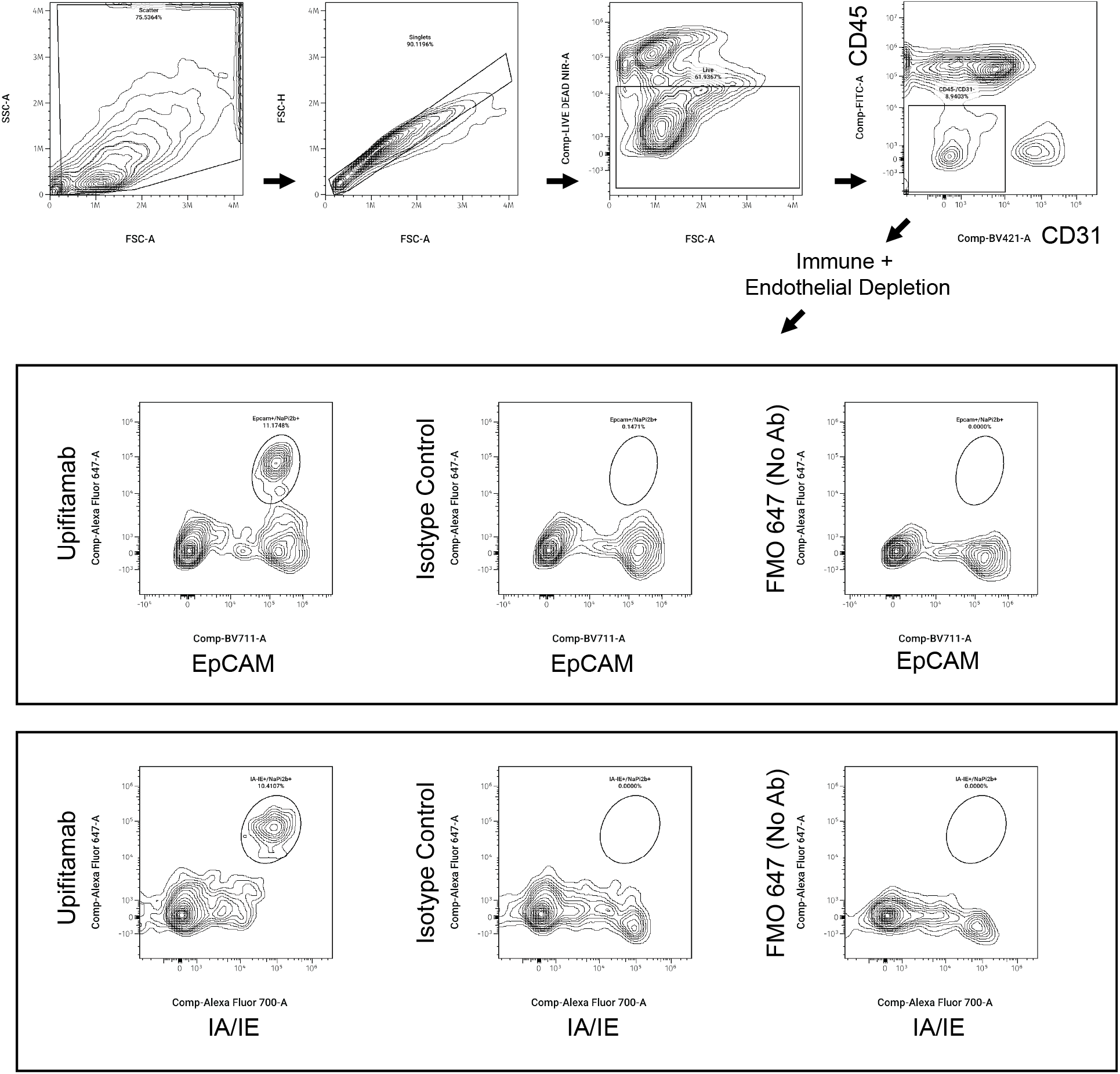
Upifitamab can be combined with IA/IE (MHC II) or EpCAM to positivity to identify mouse AT2 cells. Dissociated mouse lung was subjected to live cell staining followed by data acquisition on a Cytek Aurora 5 laser spectral flow cytometer. Following gating for singlets and viability, CD45-/CD31-cells were further analyzed by comparing expression of NaPi2b alongside EpCAM or IA/IE. Controls included an isotype control Ab conjugated to Alexa-647 and a fluorescence minus one (FMO) that contained no Alexa-647 conjugated Ab. The double-positive population (AT2 cells) is present with Upifitamab but absent with isotype IgG kappa or FMO controls. Results are representative of 3 separate experiments.

